# The coiled-coil domain of myosin-11 as a potential biomarker to detect patients with atherosclerotic cardiovascular disease risk

**DOI:** 10.1101/2025.01.28.635387

**Authors:** Lisa Takahashi, Hirofumi Tomiyama, Hiroki Kobori, Yuko Hidaka, Shunichiro Orihara, Kazuhiro Satomi, Masataka Taguri, Taishiro Chikamori, Utako Yokoyama

## Abstract

**Background:** Atherosclerosis leads to severe clinical complications, and biomarkers that could predict its progression are highly desired. Phenotypic changes and dysregulation of smooth muscle cells contribute substantially to both subclinical and clinical atherosclerosis. Plasma levels of myosin-11, a smooth muscle cell–specific myosin, have been reported to increase in patients with clinical manifestations of atherosclerosis, including coronary artery disease and/or peripheral artery disease. This study aimed to investigate the association between circulating myosin-11 levels and atherosclerotic cardiovascular disease (ASCVD) risk in patients without symptomatic atherosclerosis.

**Methods and results:** This prospective study recruited 161 participants: 80 patients with either hypertension, dyslipidemia, or diabetes mellitus but without symptomatic atherosclerosis (ASCVD risk group); 50 patients with ischemic heart disease (IHD group); and 31 control participants (Controls). We developed rat monoclonal antibodies specific for the coiled-coil domain of myosin-11 and quantified circulating myosin-11 levels using an enzyme-linked immunosorbent assay with these antibodies. Myosin-11 levels were highest in the IHD group, followed by the ASCVD risk group. Both groups had significantly higher myosin-11 levels than Controls. The area under the receiver-operating characteristic curve of myosin-11 was 0.980 for IHD versus Controls and 0.874 for ASCVD risk versus Controls. Univariate analysis detected significant associations between myosin-11 levels and hypertension and age. The association between myosin-11 levels and hypertension remained significant after multivariate logistic regression analysis.

**Conclusions:** Circulating myosin-11 levels are elevated in patients with ASCVD risk. Myosin-11 levels may be a useful biomarker for stratification of asymptomatic individuals with higher risk of ASCVD.

**Clinical Perspective:** *What Is New?:*

- We developed a sandwich enzyme-linked immunosorbent assay (ELISA) consisting of rat monoclonal antibodies specific for the coiled-coil domain of myosin-11 to detect trace concentrations of myosin-11 in blood circulation.
- Plasma myosin-11 concentrations were elevated in both asymptomatic individuals with traditional risk factors for atherosclerotic cardiovascular disease and in patients with symptomatic coronary atherosclerosis.
- This elevation may be a potential marker independent of other circulating biomarkers, traditional risk score, and non-invasive imaging analyses.

*What Are the Clinical Implications?:*

- Circulating myosin-11 may serve as a potential biomarker for identifying dysregulation of arterial smooth muscle cells during the progression of atherosclerosis.
- Further studies are warranted to evaluate whether myosin-11 levels, in combination with other reported biomarkers and risk scores, could aid in stratification of asymptomatic individuals with higher risk of ASCVD.

## INTRODUCTION

Cardiovascular disease is the leading cause of death globally, with atherosclerosis being a major underlying pathological condition responsible for cardiovascular diseases, including ischemic heart disease, peripheral artery disease, and ischemic stroke.^1–3^ The number of patients with atherosclerosis has been increasing for decades.^4^ Atherosclerosis progresses asymptomatically for an extended period before eventually leading to critical clinical events. Although numerous studies have reported associations between various circulating biomarkers and atherosclerosis,^5,6^ the detection of subclinical atherosclerosis remains a significant unmet need.^7,8^

In subclinical atherosclerosis, vascular smooth muscle cells (SMCs) undergo dysregulation.^9,10^ Rigorous SMC-specific conditional lineage tracing studies have shown that a substantial portion of cells in atherosclerotic plaque and necrotic core are derived from vascular SMCs, highlighting their importance.^9,10^ Thinning of the fibrous cap, induced by cell death and extracellular matrix breakdown, eventually leads to thrombotic events associated with the acute rupture or erosion of unstable plaques such as myocardial infarction and stroke. Additionally, evidence of multiple plaque ruptures and repairs are frequently observed, even in subclinical atherosclerosis.^9^ The presence of SMC apoptosis has also been demonstrated in early atherosclerotic lesions such as Stary/American Heart Association grades I-III, and it increases during disease progression in the necrotic core and fibrous cap.^9,11^ Based on these findings, we hypothesized that monitoring vascular SMC damage could serve as a strategy for detecting asymptomatic individuals with higher risk of atherosclerotic cardiovascular disease (ASCVD) or subclinical atherosclerosis.

Myosin-11 is specific to smooth muscle, and elevated circulating levels have been reported in patients with coronary artery disease and/or peripheral artery disease.^11^ Myosin-11 likely leaks from dead SMCs, as it is a cytoskeletal protein that is theoretically not released from viable SMCs. Myosin-11 consists of a motor domain, which contains an actin-binding site, and a coiled-coil domain. It has been demonstrated, through LC-MS/MS, that secretions from human atherosclerotic aortas mainly contained the coiled-coil domain of myosin-11.^12^ In this study, we generated monoclonal antibodies specific for the coiled-coil domain of myosin-11 to detect trace amounts of myosin-11 in blood circulation. We enrolled individuals with traditional atherosclerotic cardiovascular risk factors and patients with ischemic heart disease to investigate the association between circulating myosin-11 levels and asymptomatic individuals at risk for ASCVD.

## MATERIALS AND METHODS

### Study Subjects

From April 2020 to July 2021, 130 patients ranging in age from 50 to 79 years at Tokyo Medical University Hospital were enrolled in this study. The patients comprised two groups: an ischemic heart disease (IHD) group and a cardiovascular disease risk (ASCVD-R) group. Fifty patients who underwent catheter intervention or coronary artery bypass surgery for IHD were recruited into the IHD group. Plasma samples from the IHD group were taken within 1 week before the patients received catheter intervention or coronary artery bypass surgery. Eighty patients who received medical treatment for either hypertension, dyslipidemia, or diabetes mellitus, but who had no symptomatic atherosclerosis (including IHD, cerebrovascular disease, or peripheral artery disease) and no aortic aneurysm were classified into the ASCVD-R group. For patients in the ASCVD-R group, we confirmed, using coronary computed tomography angiography, that they did not have significant (>50%) stenosis, coronary thrombus associated with plaque rupture, or calcified nodules in coronary arteries. Plasma samples from 31 control individuals (Controls) were collected from age- and sex ratio–matched healthy volunteers at Maebashi Hirosegawa Clinic from June 2013 to January 2015. The control participants had no history of hypertension, dyslipidemia, diabetes mellitus, IHD, aortic aneurysm, cerebrovascular disease, cancer, or renal insufficiency.

Smoking status, hypertension, dyslipidemia, and the use of statins, β-blockers, angiotensin-converting enzyme (ACE) inhibitors, angiotensin receptor blockers (ARBs), calcium channel blockers (CCBs), and diuretics were documented. Body mass index, systolic and diastolic blood pressures, brachial-ankle pulse wave velocity (baPWV), and carotid intima-media thickness (IMT) were measured at the time of plasma sample collection. Plasma samples were centrifuged at 3,000 × *g* for 12 min, divided into aliquots, and stored at −80°C until analysis.

The protocol of this study was approved by the Ethical Guidelines Committee of Tokyo Medical University (reference number: T2019-0220) and adhered to the principles outlined in the Declaration of Helsinki. All human samples were obtained after written informed consent was obtained.

### Measurements of Noninvasive Physiological Parameters

Carotid IMT and baPWV were examined in the IHD and ASCVD-R groups. Ultrasonographic images were obtained with a longitudinal view of the common carotid artery. The distance between the leading edge of the lumen-intima interface and that of the media-adventitia interface was measured at multiple points along the arterial wall. The maximum IMT value was used for analysis. baPWV was measured bilaterally using a volume-plethysmographic apparatus (Form/ABI, Omron Healthcare Co., Ltd., Kyoto, Japan) with the participant supine, following the previously described method.^13^ The higher baPWV value from both sides was used for analysis.

### Generation of Anti-Myosin-11 Antibodies

The coiled-coil domain of human *MYH11* (837-1929 of accession number: NP_002465.1) was inserted into the pFLAG-CMV-3 Expression Vector (Sigma-Aldrich, St. Louis, MO, USA), and transfected in HEK293 cells. The myosin-11 fragments were purified using Toyopearl DEAE-650M and TSK-GEL G3000SW_XL_ (TOSOH Co., Tokyo, Japan) and used as recombinant antigen.

Immunization was conducted on Wistar rats (5 weeks old). Purified recombinant antigen (100 μg) was emulsified with Freund’s adjuvant (Sigma-Aldrich) and administered via footpad immunization. Immunizations were conducted every other week over a period of two months. Two month after the initial immunization, B cells were harvested from the inguinal lymph nodes of the rats. These B cells were fused with mouse myeloma cells (SP2/0) using an electrofusion method to generate hybridomas. The resulting hybridomas were cultured in E-RDF medium (Kyokuto Pharmaceutical Industrial, Kawagoe, Japan) supplemented with HAT medium supplement Hybri-Max (Sigma-Aldrich) and fetal bovine serum (FBS, Thermo Fisher Scientific). Culture supernatants from the hybridomas were screened using an enzyme-linked immunosorbent assay (ELISA) with recombinant antigen, resulting in the identification of two hybridomas producing solid-phase and detection antibodies. Western blotting was performed as described previously^14^ and demonstrated that these antibodies detected recombinant proteins of coiled-coil domain of myosin-11 (Supplemental Figure 1A).

### Myosin-11 ELISA

A sandwich ELISA using the above-mentioned rat monoclonal antibodies was performed to quantify myosin-11 levels in plasma. First, 96-well microplates were coated overnight at 4°C with the capture antibody specific to myosin-11 diluted in carbonate buffer (pH 9.6) buffer to a concentration of 2 μg/mL. Then, after coating, plates were washed with wash buffer (phosphate-buffered saline with Tween-20 [PBS-T]) to remove unbound antibodies. Non-specific binding sites were blocked with 3% bovine serum albumin for 2 h at room temperature. Plasma samples diluted 5-fold were added to the wells and incubated for 1.5 h at room temperature. Following sample incubation and washing the wells with wash buffer, the Biotin-labeled detection antibody specific to myosin-11 was added and incubated for 1 h at room temperature. After washing away the unbound detection antibody, HRP-conjugated streptavidin was added and incubated for 30 min at room temperature. After washing, HRP solution was added and incubated for 30 min at room temperature. The reaction was stopped by adding phosphoric acid to each well. Absorbance was measured at a wavelength of 450 nm using a microplate reader. The absorbance values were then correlated with the concentration of myosin-11 using a standard curve generated with known concentrations of the myosin-11 recombinant protein (Supplemental Figure 1B). The concentrations of myosin-11 in the plasma samples were determined by interpolation from the standard curve.

### High-Sensitivity C-Reactive Protein Measurement

Serum concentrations of high-sensitivity C-reactive protein (hsCRP) were measured using the N-Assay LA CRP-T kit (Nittobo Medical, Fukushima, Japan).

### Statistical Analysis

To summarize baseline characteristics and certain clinical parameters, several statistical tests were performed. For comparisons of continuous variables among three groups, the Kruskal-Wallis test was used. If statistical significance was detected, the Dunn test was conducted for pairwise group comparisons. For categorical variables, the Fisher’s exact test was applied among three groups, and if statistical significance was observed, it was also used for pairwise group comparisons. These procedures were according to Fisher’s LSD method to control the α-error in multiple comparisons. Additionally, Spearman’s correlation analysis was performed to assess relationships between clinical parameters. To evaluate diagnostic performance, receiver operating characteristic (ROC) analysis was conducted using a binary classification of factors. Cut-off values for ROC were detected using Youden index. Furthermore, univariate and multivariate linear regression analyses were performed. All statistical analyses were performed using SPSS (version 29; IBM Co., Armonk, NY, USA) and R version 4.4.2 (2024-10-31 ucrt); *P*-values <0.05 were considered statistically significant.

## RESULTS

### Characteristics of Enrolled Subjects

Clinical baseline characteristics of the ASCVD-R group, the IHD group, and Controls are shown in Table 1. There were no significant differences in age, sex distribution, or renal function among the three groups. The level of HDL-cholesterol was similar between the ASCVD-R group and the Controls and was lower in the IHD group than in the other groups. The level of LDL-cholesterol was lower in the IHD group (controlled using statins) than in the ASCVD-R group and the Controls. Triglyceride levels were higher in the IHD and ASCVD-R groups than in the Controls, with no difference between the IHD and the ASCVD-R groups. The use of statins was 100% in the IHD group and 32.5% in the ASCVD-R group. The level of hemoglobin A1c (HbA1c) was highest in the IHD group, followed by the ASCVD-R group; HbA1c levels were significantly higher in both of these groups than in the Controls. Similarly, the prevalence of diabetes mellitus was higher in the IHD group than in the ASCVD-R group. The proportion of patients with a past smoking history was greater in the IHD group than in the ASCVD-R group, with no difference between the ASCVD-R group and the Controls. Both the ASCVD-R and IHD groups had a higher frequency of hypertension compared with the Controls.

**Table 1.**
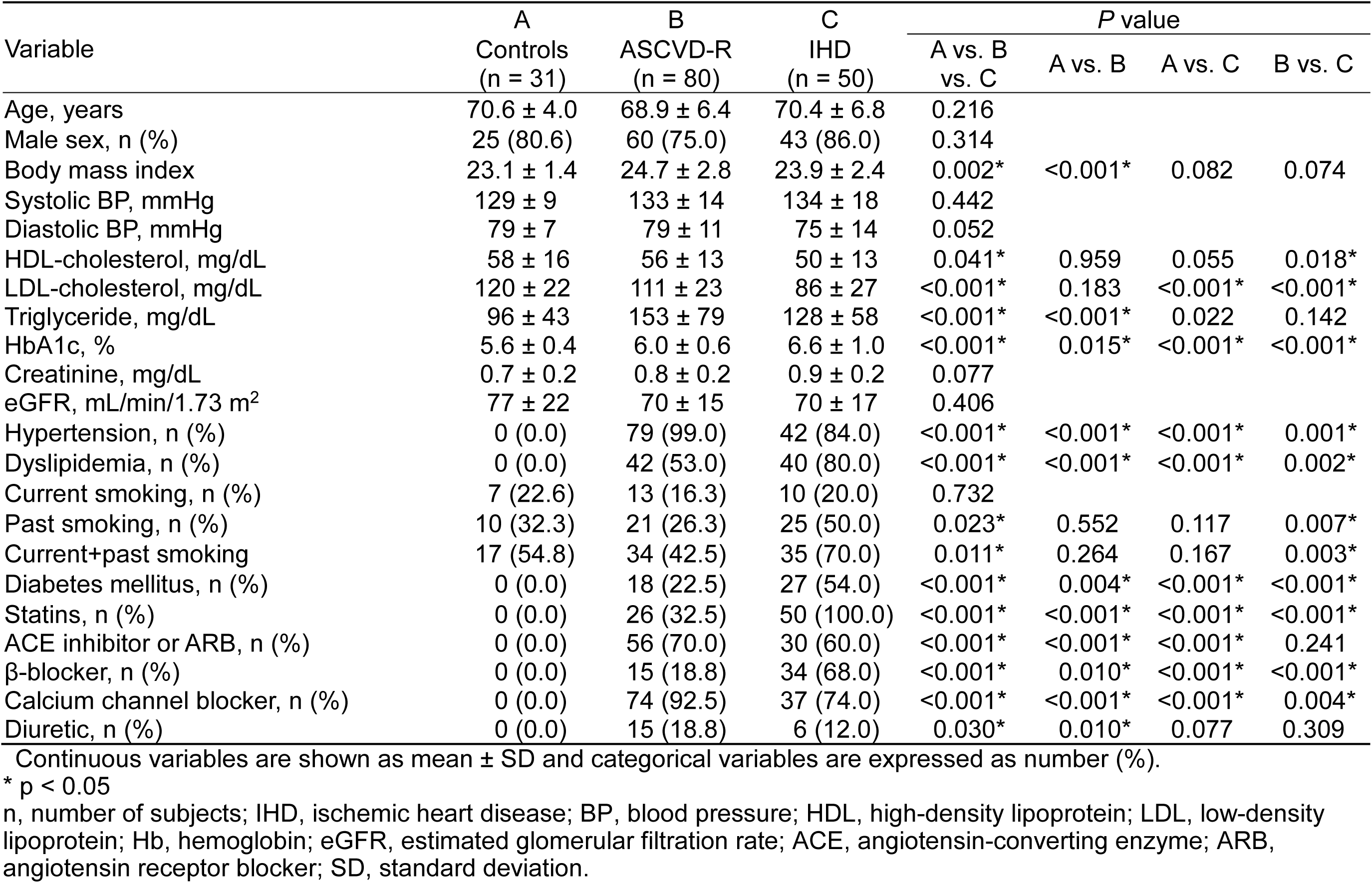
Clinical Baseline Characteristics of Patients.

| Variable | A | B | C | P value |  |  |  |
| --- | --- | --- | --- | --- | --- | --- | --- |
|  | Controls<br>(n = 31) | ASCVD-R<br>(n = 80) | IHD<br>(n = 50) | A vs. B<br>vs. C | A vs. B | A vs. C | B vs. C |
| Age, years | 70.6 ± 4.0 | 68.9 ± 6.4 | 70.4 ± 6.8 | 0.216 |  |  |  |
| Male sex, n (%) | 25 (80.6) | 60 (75.0) | 43 (86.0) | 0.314 |  |  |  |
| Body mass index | 23.1 ± 1.4 | 24.7 ± 2.8 | 23.9 ± 2.4 | 0.002* | <0.001* | 0.082 | 0.074 |
| Systolic BP, mmHg | 129 ± 9 | 133 ± 14 | 134 ± 18 | 0.442 |  |  |  |
| Diastolic BP, mmHg | 79 ± 7 | 79 ± 11 | 75 ± 14 | 0.052 |  |  |  |
| HDL-cholesterol, mg/dL | 58 ± 16 | 56 ± 13 | 50 ± 13 | 0.041* | 0.959 | 0.055 | 0.018* |
| LDL-cholesterol, mg/dL | 120 ± 22 | 111 ± 23 | 86 ± 27 | <0.001* | 0.183 | <0.001* | <0.001* |
| Triglyceride, mg/dL | 96 ± 43 | 153 ± 79 | 128 ± 58 | <0.001* | <0.001* | 0.022 | 0.142 |
| HbA1c, % | 5.6 ± 0.4 | 6.0 ± 0.6 | 6.6 ± 1.0 | <0.001* | 0.015* | <0.001* | <0.001* |
| Creatinine, mg/dL | 0.7 ± 0.2 | 0.8 ± 0.2 | 0.9 ± 0.2 | 0.077 |  |  |  |
| eGFR, mL/min/1.73 m<sup>2</sup> | 77 ± 22 | 70 ± 15 | 70 ± 17 | 0.406 |  |  |  |
| Hypertension, n (%) | 0 (0.0) | 79 (99.0) | 42 (84.0) | <0.001* | <0.001* | <0.001* | 0.001* |
| Dyslipidemia, n (%) | 0 (0.0) | 42 (53.0) | 40 (80.0) | <0.001* | <0.001* | <0.001* | 0.002* |
| Current smoking, n (%) | 7 (22.6) | 13 (16.3) | 10 (20.0) | 0.732 |  |  |  |
| Past smoking, n (%) | 10 (32.3) | 21 (26.3) | 25 (50.0) | 0.023* | 0.552 | 0.117 | 0.007* |
| Current+past smoking | 17 (54.8) | 34 (42.5) | 35 (70.0) | 0.011* | 0.264 | 0.167 | 0.003* |
| Diabetes mellitus, n (%) | 0 (0.0) | 18 (22.5) | 27 (54.0) | <0.001* | 0.004* | <0.001* | <0.001* |
| Statins, n (%) | 0 (0.0) | 26 (32.5) | 50 (100.0) | <0.001* | <0.001* | <0.001* | <0.001* |
| ACE inhibitor or ARB, n (%) | 0 (0.0) | 56 (70.0) | 30 (60.0) | <0.001* | <0.001* | <0.001* | 0.241 |
| β-blocker, n (%) | 0 (0.0) | 15 (18.8) | 34 (68.0) | <0.001* | 0.010* | <0.001* | <0.001* |
| Calcium channel blocker, n (%) | 0 (0.0) | 74 (92.5) | 37 (74.0) | <0.001* | <0.001* | <0.001* | 0.004* |
| Diuretic, n (%) | 0 (0.0) | 15 (18.8) | 6 (12.0) | 0.030* | 0.010* | 0.077 | 0.309 |

Continuous variables are shown as mean ± SD and categorical variables are expressed as number (%).
\* p < 0.05
n, number of subjects; IHD, ischemic heart disease; BP, blood pressure; HDL, high-density lipoprotein; LDL, low-density lipoprotein; Hb, hemoglobin; eGFR, estimated glomerular filtration rate; ACE, angiotensin-converting enzyme; ARB, angiotensin receptor blocker; SD, standard deviation.

The use of ACE inhibitors, ARBs, and diuretics was similar between the IHD and ASCVD-R groups. The use of β-blockers was higher in the IHD group than in the ASCVD-R group, and the use of calcium channel blockers was higher in the ASCVD-R group than in the IHD group. None of the Controls used statins, ACE inhibitors, ARBs, β-blockers, calcium channel blockers, or diuretics.

### Plasma Myosin-11 Levels

We measured plasma concentrations of myosin-11 in the three groups using an ELISA specific for the coiled-coil domain of myosin-11. Plasma myosin-11 levels were significantly higher in the IHD group (median: 0.922 μg/mL) than in the Controls (median: 0.062 μg/mL), which is consistent with the previous reports.^11^ Myosin-11 levels in the ASCVD-R group (median: 0.495 μg/mL) were lower than in the IHD group but significantly higher than in the Controls (Figure 1A).

**Figure 1.**
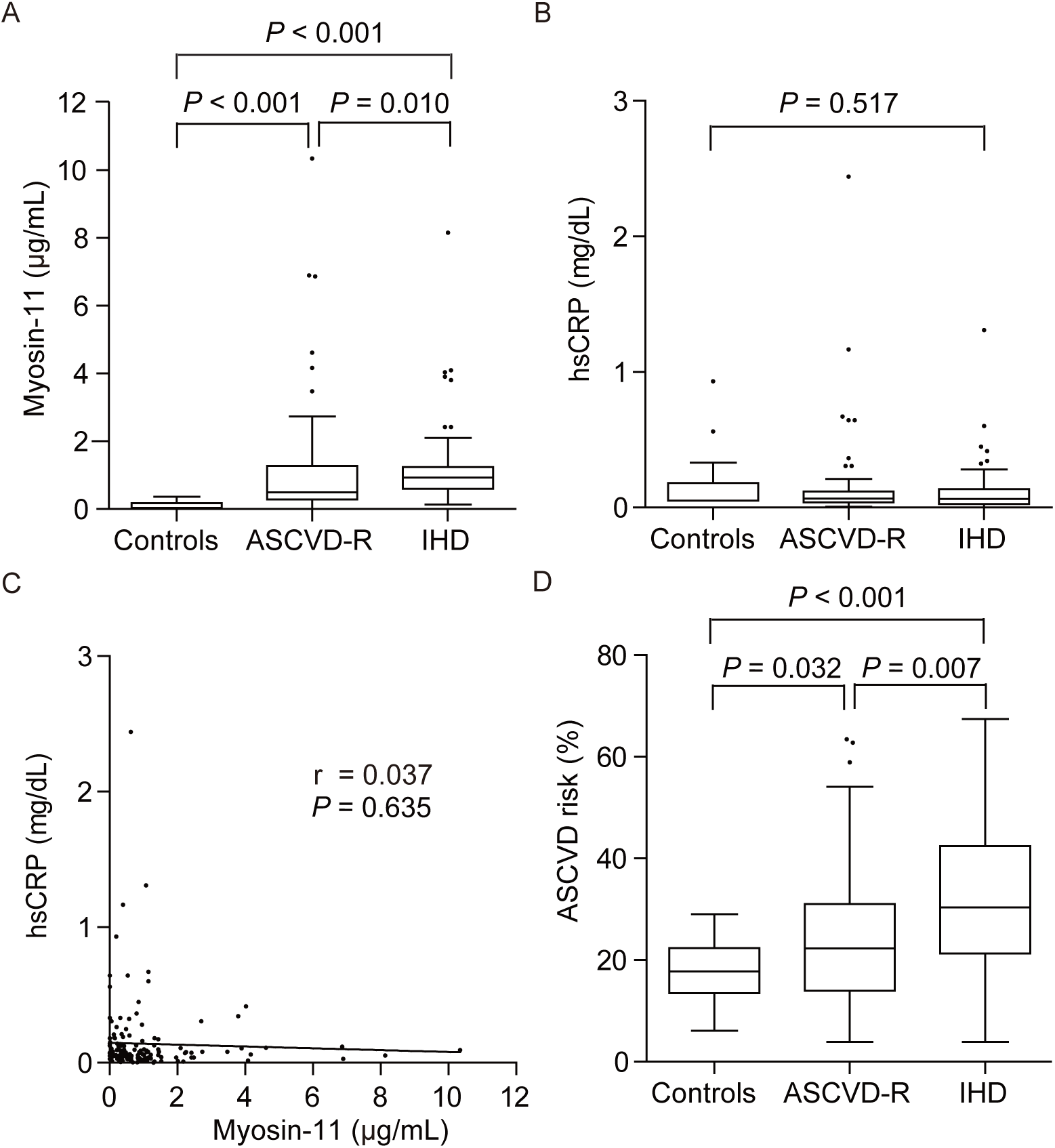
Comparison of plasma myosin-11 levels, hsCRP, and 10-year ASCVD risk score in the ASCVD-R, IHD, and Control groups. **A.** Myosin-11 concentrations in plasma samples of controls (Controls, n=31), patients with ASCVD risk factors (ASCVD-R, n=80), and patients with ischemic heart disease (IHD, n=50). **B.** hsCRP concentrations in plasma samples in the 3 groups. **C.** Correlation between plasma concentrations of myosin-11 and hsCRP in all participants (*n*=161). **D.** Ten-year ASCVD risk score in the 3 groups. Data of A, B, and D are shown as Tukey boxplots.

In this sample set, we analyzed serum concentrations of hsCRP, which is the most extensively discussed potential biomarker for atherosclerosis.^15,16^ Concentrations of hsCRP did not differ significantly among the three groups (Figure 1B). No correlation was observed between the concentrations of myosin-11 and hsCRP (Figure 1C). These data suggest that the concentrations of myosin-11, as detected by this ELISA, were associated not only with the presence of symptomatic coronary atherosclerosis but also with the accumulation of cardiovascular risk factors.

We also calculated the ASCVD risk score using 2018 Prevention Guidelines Tool ASCVD-Risk Calculator (https://static.heart.org/riskcalc/app/index.html#!/baseline-risk), which is used to estimate an individual’s 10-year ASCVD risk based on age, sex, race, total cholesterol, HDL-cholesterol, systolic blood pressure, blood pressure–lowering medication use, diabetes status, and smoking status.^17,18^ The scores were significantly higher in the IHD group than in the Controls (Figure 1D). The scores of the ASCVD-R group were lower than in the IHD group but significantly higher than in the Controls, as expected.

These data suggest that this set of monoclonal antibodies, specific to the coiled-coil domain of myosin-11, has the potential to discriminate both symptomatic and asymptomatic individuals with ASCVD risks from control subjects.

### Efficacy of Myosin-11 for Diagnosis of Patients with IHD and ASCVD Risks

We evaluated the diagnostic value of myosin-11 using ROC analysis of myosin-11 to detect the accumulation of ASCVD risk or the presence of atherosclerosis. When comparing the Controls with the ASCVD-R group, the area under the curve (AUC) of myosin-11 was 0.874 (95% CI: 0.810-0.937), with a specificity of 96.6% at a sensitivity of 72.5%. The positive predictive value, negative predictive value, accuracy, and cutoff value were 98.3%, 56.0%, 78.9%, and 0.322 μg/mL, respectively. The AUC of hsCRP was 0.467 (95% CI: 0.344-0.589), with a specificity of 64.6% at a sensitivity of 58.1%; positive predictive value, negative predictive value, accuracy, and cutoff value were 78.3%, 36.0%, 59.1%, and 0.053 mg/dL, respectively. The AUC of ASCVD risk was 0.640 (95% CI: 0.535-0.745), with a specificity of 48.8% at a sensitivity of 80.6%; positive predictive value, negative predictive value, accuracy, and cutoff value were 86.7%, 37.9%, 57.7%, and 22.5%, respectively (Figure 2A). The AUC of myosin-11 was significantly greater than that of hsCRP and ASCVD risk (*P*<0.001 and *P*<0.001, respectively).

**Figure 2.**
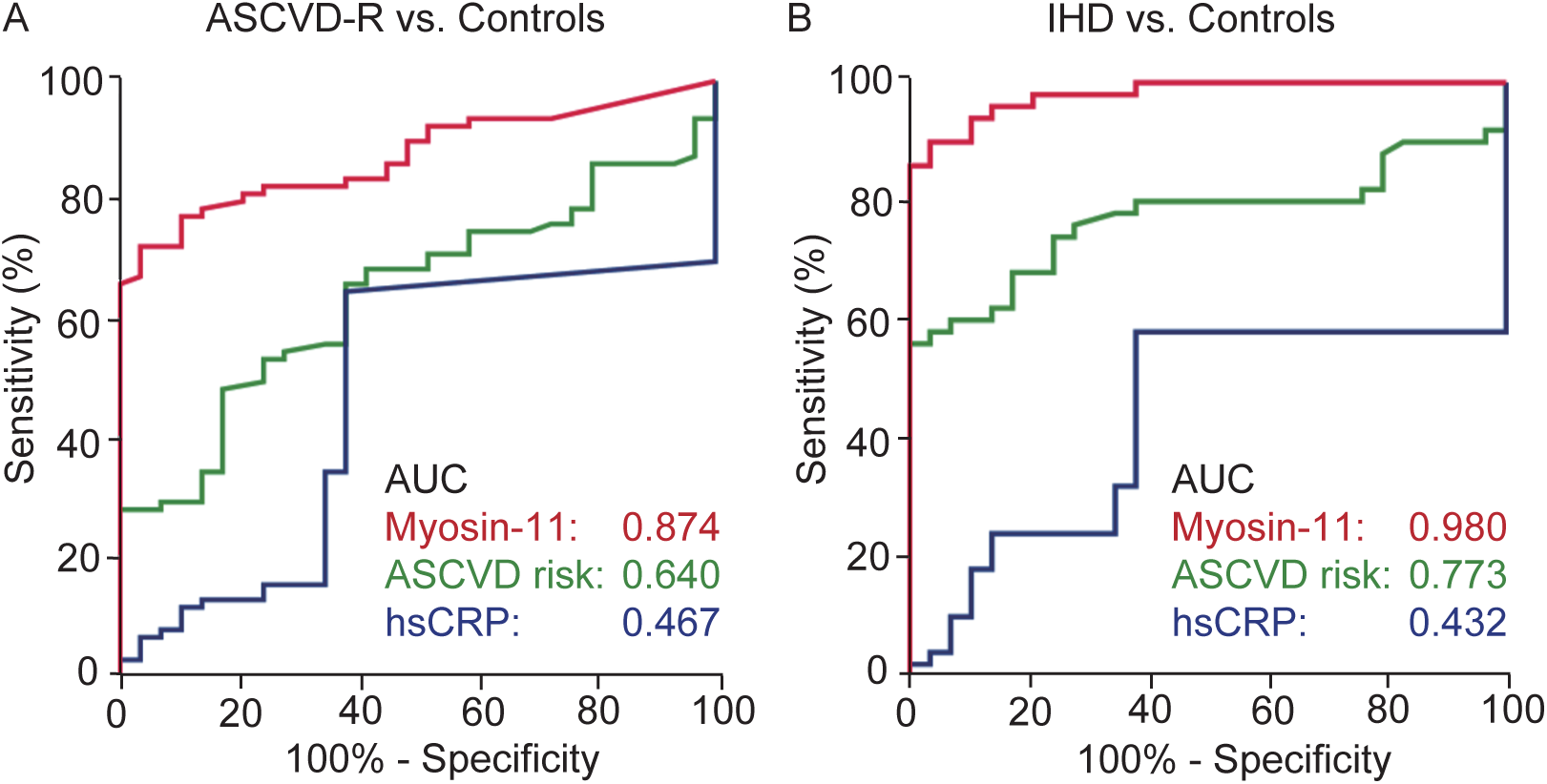
ROC analysis of myosin-11 and hsCRP. **A.** ROC curve of myosin-11, ASCVD risk, and hsCRP in the ASCVD-R group and Controls. **B.** ROC curve of myosin-11, ASCVD risk, and hsCRP in the IHD group and Controls. Controls, n=31; ASCVD-R, n=80; IHD, n=50.

When comparing the Controls with the IHD group, the AUC of myosin-11 was 0.980 (95% CI: 0.957-1.003), with a specificity of 90.0% at a sensitivity of 96.6%; positive predictive value, negative predictive value, accuracy, and cutoff value were 97.8%, 84.8%, 92.4%, and 0.326 μg/mL, respectively. The AUC of hsCRP was 0.432 (95% CI: 0.304-0.561), with a specificity of 58.0% at a sensitivity of 52.0%; positive predictive value, negative predictive value, accuracy, and cutoff value were 69.0%, 46.2%, 84.0%, and 0.052 mg/dL, respectively. The AUC of ASCVD risk was 0.773 (95% CI: 0.669-0.877), with a specificity of 56.0% at a sensitivity of 100%; positive predictive value, negative predictive value, accuracy, and cutoff value were 100%, 58.5%, 72.8%, and 22.5%, respectively (Figure 2B). The AUC of myosin-11 was significantly greater than that of hsCRP and ASCVD risk (*P*<0.001 and *P*<0.001, respectively).

### Association of Myosin-11 Concentrations and Physiological Parameters of Atherosclerosis

Carotid IMT is widely recognized as an early surrogate marker of subclinical atherosclerosis and a strong predictor of future cardiovascular events.^19–22^ In this sample set, the values of maximum carotid IMT were significantly greater in the IHD group (median: 2.95 mm) than in the ASCVD-R group (median: 1.90 mm) (Figure 3A). However, there was no correlation between myosin-11 concentrations and maximum carotid IMT (Figure 3B). Arterial stiffness is associated with progression of atherosclerosis, and elevated baPWV is an independent predictor of the presence of obstructive coronary artery disease.^23,24^ Both the IHD (median: 1709 cm/s) and ASCVD-R (median: 1707 cm/s) groups had similar and higher values, which is accordance with previous reports showing that baPWV value ranges from 1500 to 1800 cm/s in patients with cardiovascular disease or its risk factors (Figure 3C).^23^ Similarly to the carotid IMT results, baPWV was not correlated with myosin-11 concentrations (Figure 3D). These data suggest that myosin-11 levels are independent of these noninvasive physiological parameters.

**Figure 3.**
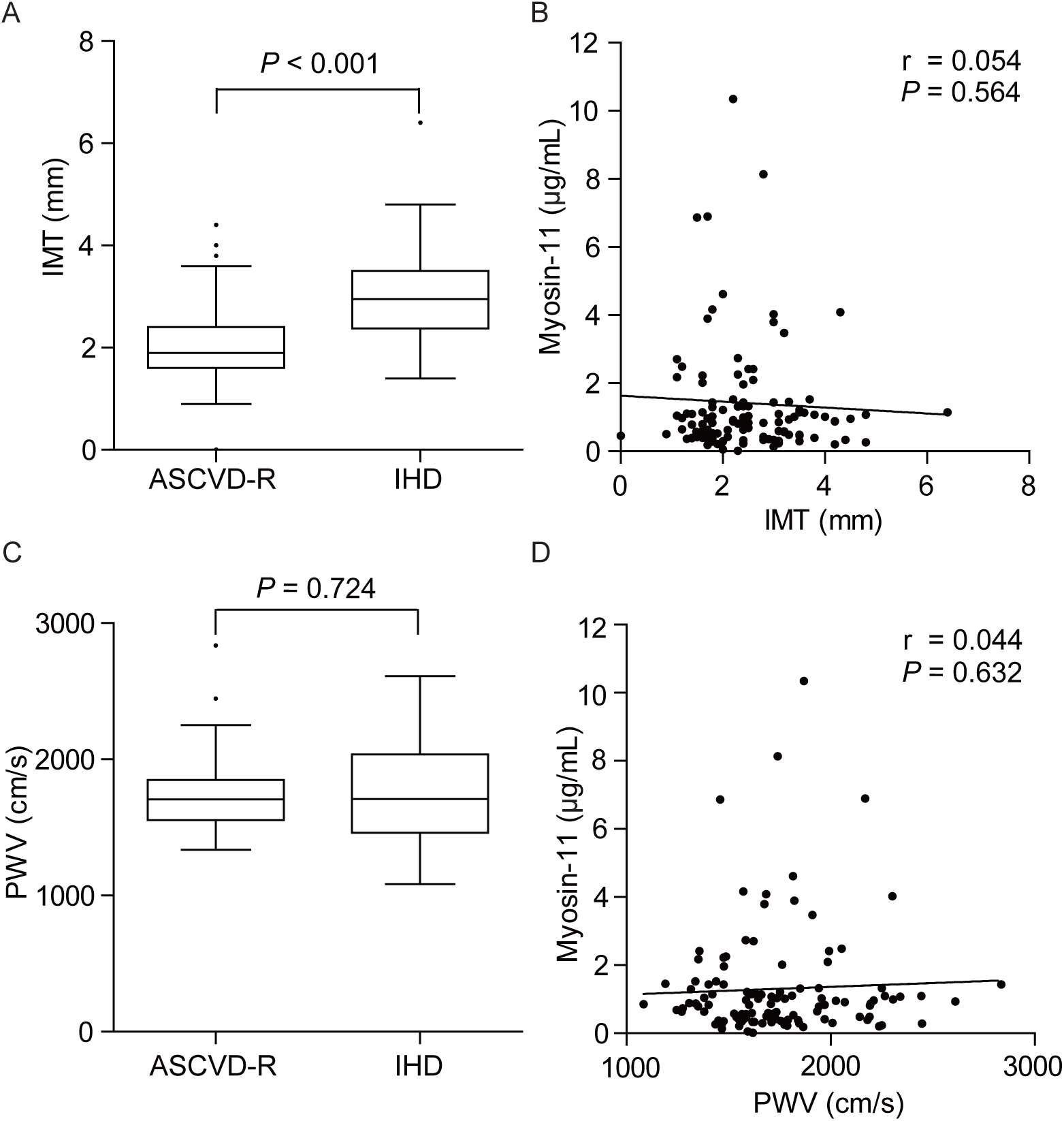
Association of myosin-11 levels and physiological parameters of atherosclerosis. **A and C.** Comparison of IMT and baPWV between the ASCVD-R and IHD groups. **B and D.** Correlation between plasma concentration of myosin-11 and each of IMT and baPWV in the ASCVD-R and IHD groups. ASCVD-R, n=80; IHD, n=50. Data of A and C are shown as Tukey boxplots.

### Association Between Circulating Myosin-11 and Risk Factors for ASCVD

We further investigated which traditional risk factors were associated with elevating circulating myosin-11 concentrations. Among the risk factors for ASCVD, including hypertension, dyslipidemia, diabetes mellitus, and smoking history, univariate analysis demonstrated a significant association between myosin-11 levels and both age and hypertension (Table 2). This significant association between increased myosin-11 levels and hypertension persisted in the multivariate analysis (Table 3).

**Table 2.** Univariate Analysis Between Risk Factors for Atherosclerosis and Plasma Concentration of Myosin-11 in all Participants.

| | $\beta$ | *P* value | 95% CI |
| --- | --- | --- | --- |
| Age | -0.172 | 0.030 | -0.078– -0.004 |
| Male sex | -0.005 | 0.945 | -0.595– 0.555 |
| Body mass index | 0.127 | 0.110 | -0.017– 0.166 |
| Hypertension | 0.295 | <0.001 | 0.499– 1.543 |
| Dyslipidemia | 0.153 | 0.055 | -0.009– 0.912 |
| Smoking history | -0.089 | 0.264 | -0.728– 0.201 |
| Diabetes mellitus | 0.110 | 0.168 | -0.154– 0.875 |

**Table 3.** Multivariate Analysis Between Risk Factors for Atherosclerosis and Plasma Concentration of Myosin-11 in all Participants.

| | $\beta$ | *P* value | 95% CI |
| --- | --- | --- | --- |
| Age | -0.149 | 0.075 | -0.076–0.004 |
| Male sex | 0.006 | 0.941 | -0.577–0.622 |
| Body mass index | 0.018 | 0.828 | -0.084–0.105 |
| Hypertension | 0.248 | 0.004 | 0.272–1.432 |
| Dyslipidemia | 0.047 | 0.582 | -0.359–0.638 |
| Smoking history | -0.078 | 0.351 | -0.717–0.256 |
| Diabetes mellitus | 0.058 | 0.504 | -0.370–0.749 |

## DISCUSSION

In this study, we developed an ELISA using rat monoclonal antibodies targeting the coiled-coil domain of myosin-11. Consistent with a previous report utilizing a commercially available ELISA based on polyclonal antibodies against myosin-11,^11^ the levels of circulating myosin-11 were significantly higher in the IHD group compared with the Controls. Additionally, our newly developed ELISA distinguished asymptomatic individuals with ASCVD risks from the Controls, with myosin-11 levels being lower in both groups than in the IHD group. Whereas traditional risk factors such as hypertension, dyslipidemia, diabetes, and smoking are well established, a significant portion of individuals with subclinical atherosclerosis may not be detected through conventional risk assessment tools.^25^ In our sample, patients’ 10-year ASCVD risk scores showed a similar trend to myosin-11 levels among the three groups, but the AUC of myosin-11 levels was greater than that of ASCVD risk scores between the ASCVD-R group and the Controls. As our investigation is cross-sectional, it remains unclear whether stratification based on myosin-11 levels between the Controls and the ASCVD-R group correlates with future cardiovascular events. A longitudinal study in the general population would be valuable to assess whether circulating myosin-11 concentration is associated with atherosclerosis progression and subsequent cardiovascular events.

The ASCVD-R group was defined as asymptomatic individuals without coronary artery lesions with >50% stenosis and no plaque-induced thrombus formation, and with at least one traditional ASCVD risk factor. This ASCVD group exhibited a variable degree of maximum carotid IMT, which is well recognized as an indicator for the presence of subclinical atherosclerosis, as shown in Figure 3A.^26^ Although there is no strict pathological definition of subclinical atherosclerosis,^27^ it is plausible that the ASCVD-R group includes both individuals at higher risk of developing atherosclerosis and those with subclinical atherosclerosis who have not yet experienced clinical events. Although additional research is needed, our findings suggest that circulating myosin-11 concentration has the potential to detect dysregulation of vascular SMCs undetectable by imaging modalities or subclinical atherosclerosis.

Numerous biomarkers have been reported in association with the progression of atherosclerosis over the past decades.^28,29^ ASCVD is characterized by a complex pathological process involving multiple disease stages,^30–32^ with biomarkers such as proteins related to lipid, inflammation, endothelial dysfunction, and coagulation being proposed. Among these, hsCRP has shown considerable promise in predicting future cardiovascular events, such as myocardial infarction and ischemic stroke.^33,34^ However, nearly half of patients experiencing acute coronary syndrome have low or very low levels of CRP, highlighting residual ASCVD risks unrelated to persistent sterile inflammation.^3^ In this study, hsCRP levels did not differ significantly among the ASCVD-R, IHD, and Control groups, and were not associated with myosin-11 concentration. Myosin-11 is a cytoskeletal protein primarily expressed in arterial SMCs, and circulating myosin-11 is believed to reflect leakage from damaged vascular SMCs, under conditions with or without inflammation. Therefore, detection of circulating myosin-11 may also capture non-inflammatory mechanisms of atherosclerosis.

Extracellular vesicles represent a heterogeneous population of vesicles secreted by various cell types during atherosclerosis progression, including immune responses, cell proliferation, migration, and cell death.^35^ MicroRNAs (miRs) carried by extracellular vesicles have been widely studied for their association with atherosclerosis, with several SMC-specific miRs proposed as potential biomarkers.^35–38^ For example, circulating miR-221 levels were significantly lower in patients with coronary stenosis,^36^ and miR-143 was inversely associated with time-to-coronary artery disease incident event.^38^ Although myosin-11 is not thought to be secreted from cells under normal conditions, it reportedly has two major protease-susceptible sites located at the head-rod junction and the heavy meromyosin-light meromyosin junction.^39^ Tryptic and chymotryptic digestion produces light meromyosin (80-85 kDa) and short subfragment 2 (40-45 kDa) segments.^39^ Based on these findings, it is possible that proteolytic cleavage of myosin-11 produces coiled-coil domain fragments, which is the opposite site of actin binding, in dysregulation of SMCs and that these fragments can be transferred via extracellular vesicles from viable SMCs to the blood circulation. It has been reported that myosin-9, which has a similar structure to myosin-11, was detected in the blood circulation of Woody breast myopathy, but it remains unknown which part of myosin-9 was detected in plasma.^40^ Further studies are needed to elucidate molecular mechanisms underlying the transfer of myosin-11 fragments from SMCs to blood circulation.

Given the complexity of atherosclerosis pathology, combining biomarkers have been shown to enhance the prediction of cardiovascular outcomes. For instance, use of the biomarkers troponin I, N-terminal pro–brain natriuretic peptide, cystatin C, and hsCRP, along with traditional risk factors (age, hypertension, dyslipidemia, diabetes mellitus, smoking history, and BMI), improves prediction of cardiovascular mortality.^41^ Atherogenic lipoproteins, such as apolipoprotein B and lipoprotein(a), are strongly associated with ASCVD risk,^42–44^ and their combination with hsCRP has proven effective in predicting cardiovascular events during a 5-year follow-up.^45^ Additionally, artificial intelligence is increasingly recognized as a valuable tool for detection of subclinical atherosclerosis and ASCVD risk stratification.^46^ Integrating multimodal data from various sources, including biomarkers detected in blood circulation, imaging, and genetic information, could better identify high-risk individuals. In such a prediction model, it would be useful to include multiple distinct pathological mechanism–related biomarkers, such as lipoproteins, inflammation, and vascular SMC damage. Our findings suggest that myosin-11 is independent of the values of hsCRP, carotid maximum IMT, and baPWV, although carotid maximum IMT and baPWV were not examined in the Control group. Myosin-11 would provide additive value to these well-recognized indicators. Further studies are needed to evaluate the utility of combining myosin-11 with other potential circulating biomarkers, traditional risk score, and non-invasive imaging analyses.

This study has several limitations. First, the sample size was small. Second, the ASCVD-R group primarily consisted of patients with hypertension. Although multivariate analysis indicated an association between increased circulating myosin-11 levels and hypertension, selection bias may have influenced this relationship. Further studies should include a larger cohort with a diverse range of risk factors. Third, this study focused on an elderly population (average age of approximately 70 years). Given that atherosclerosis is documented in youth and early adulthood,^7^ future research should evaluate the utility of circulating myosin-11 in predicting ASCVD in younger populations.

In conclusion, we developed an ELISA using monoclonal antibodies to detect trace levels of circulating myosin-11. Plasma levels of the coiled-coil domain of myosin-11 were elevated in patients with ASCVD risks and IHD compared to controls. Circulating myosin-11 may be a potential biomarker for detecting individuals with higher ASCVD risk and clinical atherosclerosis.

## Data Availability Statement

The data that support the findings of this study are available upon request from the corresponding author.

## Acknowledgments

The authors are grateful to Yuka Sawada for technical assistance.

## Nonstandard Abbreviations

ACE: angiotensin-converting enzyme
ARBs: angiotensin receptor blockers
ASCVD: atherosclerotic cardiovascular disease
AUC: area under curve
baPWV: brachial-ankle pulse wave velocity
CCBs: calcium channel blockers
HRP: horseradish peroxidase
hsCRP: high-sensitivity C-reactive protein
IHD: ischemic heart disease
IMT: intima-media thickness
ROC: receiver-operating characteristic

